# Assessing the utility of minority variant composition in elucidating RSV transmission pathways

**DOI:** 10.1101/411512

**Authors:** George Githinji, Charles N. Agoti, Nelson Kibinge, Sonal Henson, Patrick Munywoki, Samuel Brand, Graham Medley, Patricia Cane, Matthew Cotten, D. James Nokes, Colin J. Worby

## Abstract

Reconstructing transmission pathways and defining the underlying determinants of virus diversity is critical for developing effective control measures. Whole genome consensus sequences represent the dominant virus subtype which does not provide sufficient information to resolve transmission events for rapidly spreading viruses with overlapping generations. We explored whether the within-host diversity of respiratory syncytial virus quantified from deep sequence data provides additional resolution to inform on who acquires infection from whom based on shared minor variants in samples that comprised epidemiological clusters and that shared similar genetic background. We report that RSV-A infections are characterized by low frequency diversity that occurs across the genome. Shared minor variant patterns alone, were insufficient to elucidate transmission chains within household members. However, they provided inference on potential transmission links where phylogenetic methods were uninformative of transmission when consensus sequences were identical. Interpretation of minor variant patterns was tractable only for small household outbreaks.

## Introduction

Respiratory syncytial virus (RSV) is an important cause of respiratory illnesses among infants and children under the age of five years (Berkley et al. 2010; Henderson, Collier, and Clyde Jr 1979). RSV spreads rapidly within communities, usually in seasonally occurring outbreaks, and is efficiently transmitted within households, exhibiting short doubling times and infecting a high proportion of the household (Hall et al. 1981; Munywoki et al. 2014) many of whom are reinfections (Hall et al. 1991) given that immunity to infection is short-lived (Ohuma et al. 2012). RSV-associated pneumonia and subsequent mortality contributes to a substantial burden of disease in developing countries and it is estimated that 45% of pneumonia-associated hospital admissions are attributable to RSV. Infants, particularly in the first 6 months of life, are at heightened risk of disease following RSV infection (Shi et al. 2017) and are a key target for vaccine protection. Direct protection of these infants by vaccines presents a major challenge and alternative options include interruption of chains of transmission by targeting those most likely to infect the infants. For this reason, improved understanding of who acquire infection from whom in the community and particularly in the household is critical.

Understanding of how respiratory viruses spread and evolve within a population improves and informs our capacity to control outbreaks. This is greatly aided by the collection of detailed individual-level data, with which we may attempt to identify inter-host transmission pathways which is challenging for highly transmissible viruses, such as RSV, that result in explosive seasonal outbreaks, leading to overlapping generation of cases. As such, teasing out transmission pathways cannot rely on temporal case data alone and is increasingly supported by use of virus genome sequence data to establish the relationships between infecting viruses and hence who infects whom.

It is not always possible to target interventions for those most at risk of severe disease. However, outbreaks may be mitigated by preventing infection in individuals that are most important in community transmission or by selecting target individuals known to be important in infecting the at-risk individuals, for example infants, through family cocooning. This is exemplified by RSV, for which development of a vaccine to directly protect the vulnerable infant has proved fruitless to date, hence a need to explore indirect methods of protection.

RSV is characterised by substantial genetic diversity, typical of RNA viruses which are described to exist as a quasi-species (Domingo, Sheldon, and Perales 2012; Lauring and Andino 2010). Overall, within and between host variation has been suggested to impact and affect both epidemiological and clinical factors of an infection (Alizon, Luciani, and Regoes 2011). For some specific viruses, intra-host diversity has been shown to play a role in virulence (Töpfer et al. 2013), pathology and disease progression (Vignuzzi et al. 2006), immune escape (Henn et al., 2012) and may provide insight on transmission patterns (Gire et al. 2014; Paterson et al. 2015; Poon et al. 2016; Stack et al. 2013). In addition, diversity may be important for adaptation (Vignuzzi et al. 2006) and survival (Crotty, Cameron, and Andino 2001) which has a direct consequence on infectivity, fitness and pathogenesis. While one can generate consensus sequence data and apply commonly-used genomic approaches to such rapidly diversifying pathogens, it is likely that some degree of resolution will be lost by doing so. Furthermore, removing resolution may be particularly disadvantageous for rapidly spreading pathogens, where temporal information may be less informative regarding transmission routes. It is against this background that the minor variants characteristics of viruses might provide additional transmission resolution through characterisation of shared low frequency diversity between individuals.

Obtaining a detailed knowledge of infection patterns for rapidly diversifying viruses in absence of host contact and mobility data during an outbreak is a challenge. In this study we utilise data collected during a six-month community surveillance of 47 households on the coast of Kenya spanning a complete seasonal outbreak of RSV (Agoti et al. 2017; Agoti, Otieno, Ngama, et al. 2015; Munywoki et al. 2014). Nasopharyngeal samples from 9 of 47 study households were sequenced using a paired-end Illumina MiSeq protocol (Agoti, Otieno, Munywoki, et al. 2015). Using these data, we characterized the within and between host genetic diversity and explore the application of shared minority variants diversity to infer the transmission patterns of RSV from this community grouping of households. We discuss the potential and pitfalls of applying these methods to complement phylogenetic methods using consensus sequence data.

Circulating consensus RSV diversity in Kilifi has been described and it is discernible both at the community (Agoti, Otieno, Munywoki, et al. 2015) and household levels (Agoti et al. 2017). A detailed description of the entire spectrum of low frequency diversity in this set of samples has not been carried out.

## Materials and Methods

### Study population

The samples were collected as part of a study conducted in the northern part of Kilifi County and within the Kilifi Health and Demographic Surveillance System (KHDSS) (Scott et al. 2012). Households and all consenting occupants were followed for a full RSV epidemic from December 2009 to June 2010. A household was defined as a unit comprised of individuals who prepared and shared meals together (Munywoki et al. 2014; Agoti, Otieno, Munywoki, et al. 2015). The epidemiological study was conducted across 47 households with an infant (<12 months of age) and at least one elder sibling (<15 years of age). Nasopharyngeal swabs were collected from the consenting household members twice weekly regardless of symptoms as previously described and RSV positives identified through molecular diagnostic screening (Munywoki et al. 2014). The current analysis is limited to 137 RSV positive samples from nine households for which whole genome next-generation sequencing (NGS; also called deep or high-throughput sequencing) was undertaken (Agoti, Otieno, Munywoki, et al. 2015; Agoti et al. 2017).

### Bioinformatics analysis

We developed a bioinformatics pipeline to identify variant positions from short read whole genome sequence data (supplementary figure 1). Raw reads were quality trimmed and duplicated reads filtered out. Quality trimmed reads from each sample were mapped against a *de novo* assembled RSV genome and the resulting BAM files were used as input for variant calling using LoFreq (Wilm et al. 2012), Vardict (Lai et al. 2016), FreeBayes (Garrison and Marth 2012), SNVer (Wei et al. 2011) and Varscan2 (Koboldt et al. 2012).

#### Quality control

Raw FASTQ reads were processed as follows, duplicated reads were removed using QUASR (Watson et al. 2013) version 7.05 followed by quality trimmed using Trimmomatic (Bolger, Lohse, and Usadel 2014) based on a four-base sliding window which scanned the read from the 5′ end of the read, and removed the 3′ end of the read when the average quality of a group of bases dropped below a Phred score of 25 ensuring that only consecutive series of poor-quality bases triggered trimming.

#### Reference genome sequence

The reference sequence was derived from a previously assembled sequence of a sample collected from an individual in household 40, accession number KX510245.1 (Agoti et al. 2017).

#### Short read alignment

The reads from each RSV-A positive sample were aligned against the reference sequence (GenBank accession number KX510245.1) using BWA (Li 2010) version 0.7.13-r1126. The output BAM file was sorted and indexed using SamTools (Li et al. 2009) version 1.3.1.

#### Consensus sequence alignment

Consensus sequences (Agoti et al. 2017) from each sample were aligned using MAFFT (Katoh and Standley 2013) version 7.271. The resulting alignment was provided as input to IQ-TREE version 1.5.5 to infer a maximum-likelihood tree (Nguyen, Schmidt, Haeseler, & Minh, 2015). Model selection was performed automatically using the model-test functionality available from IQ-TREE.

#### Variant calling

BAM files that were generated from the short-read alignment step were used to call minority variants using five variant calling programs: LoFreq (Wilm et al. 2012) version 2.1.2, FreeBayes (Garrison and Marth 2012) version 1.1.0-3-g961e5f3, SNVer (Wei et al. 2011) version 0.5.3, VarDict (Lai et al. 2016) version 30.3.17 and VarScan (Koboldt et al. 2012) version 2.4.2. Variant frequencies were average across the all frequencies from the concordant callers and at each position.

#### Filtering variants

We only considered minor variants that were identified at positions with at least a read coverage of 200 and reported by three or more minority variant callers. Minor variants with frequencies below 1% were discarded as well as minor variants whose positions coincided with primer regions.

### Quantifying within host minority variant diversity

Within host diversity was quantified using Shannon’s entropy (Shannon 1948) that was calculated based on average frequencies of each minority variant detected by the concordant callers. Shannon’s entropy was calculated for each sample and gene region.

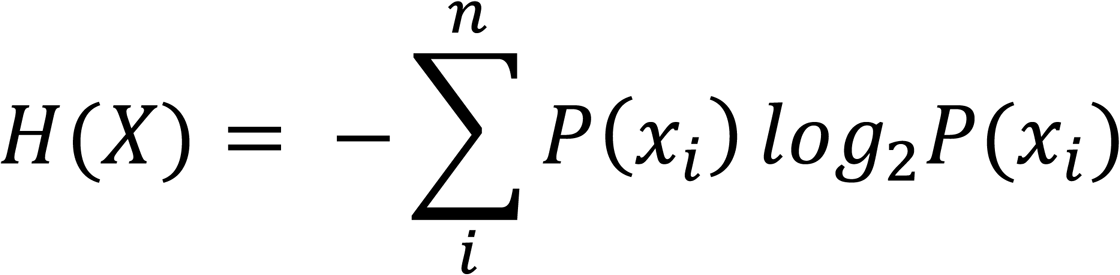

Where, *x*_*i*_ is the mean frequency (averaged over the concordant variant callers) of the minority variant at position *i* and summing over all *n* positions in the genome. To quantify temporal trend in within-host diversity, Shannon’s entropy diversity measurement was plotted against the time course profile of individuals with samples collected at multiple time points.

### Quantifying the genetic distance between samples

The L1-norm (Manhattan) distance over nucleotide frequency at each position was used to quantify within and between sample pairwise genetic distances. For each pair of samples x and y, the Manhattan distance d_i_ between two samples *x* and y d_xy_ calculated at each nucleotide position *i* is given by;

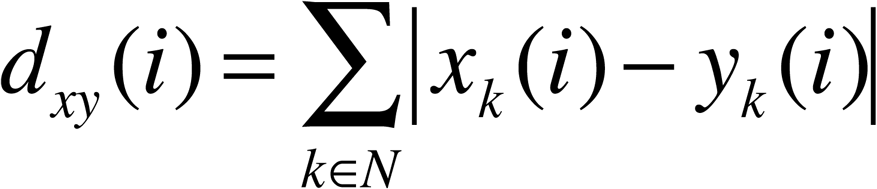

where N is the set of nucleotides *A, C, G, T*, and *x*_*k*_*(i)* is the frequency of nucleotide *k* in position *i* from sample *x*. The Manhattan distance between two samples *D*_*xy*_ is the sum of distances *d*_*xy*_ at each position in the genome.

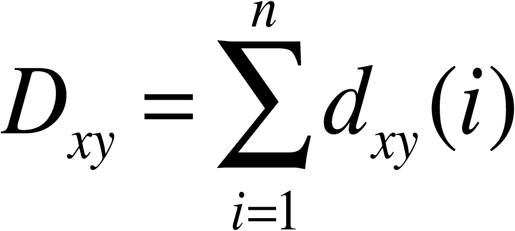

Where D is a similarity pairwise distance metric for any two samples. The samples were stratified, and the distances plotted based on whether the comparisons were within or between household memberships.

To assess the significance of differences in typical L1-norm distances observed within and between household groups, we estimated the difference in mean and standard deviation between the two groups as parameters using a regression model implemented in a Bayesian context to capture the uncertainty in mean and standard deviation given a skewed distribution.

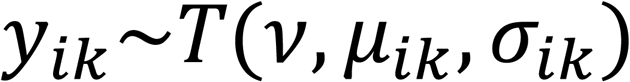

Where *i* is the *i*^*th*^ datum and *k* indicates the *k*^*th*^ group. *y*_*ik*_ are draws from a t-distribution with normality parameter *v*, mean *μ*_*ik*_ and standard deviation *<u>σ_ik_</u>* for datum *i* in group *k* We fitted this model using a Bayesian HMC approach using the BRMS package (Bürkner 2017) and Stan (Carpenter et al. 2016). By assuming a t-distribution we minimized the effect of the outliers while estimating the mean and standard deviation of the genetic distances.

### Transmission tree model using shared variants

A conservative minority variants frequency was used to identify variant positions in each sample, and we discarded variants shared by more than 10 hosts, which appeared empirically to be outliers, and would be uninformative of transmission patterns. From samples *x*_*1*_,…, *x*_*n*_ collected from individuals *1,…, n*, we identified the set of variant loci *V*_*i*_ for the *i-th* sample such that a nucleotide was observed at a frequency in the interval (0.01 <= x < 0.5) at these positions. For each pair of samples, we counted the number of shared variant positions, forming an adjacency matrix which was transformed into a weighted variant transmission tree (Worby, Lipsitch, & Hanage, 2017). We discarded variant positions that were shared by more than 10 individuals, as well as samples with more than 20 variant positions. We described the shared variants at the sample and individual (host) levels. In the latter case, any shared variants observed between any of the samples collected from two individuals were considered an inter-host link.

## Results

### Intra-host diversity of RSV-A viruses

We assessed the distribution of minority variants across all RSV-A samples at different levels of concordance (Figure 1A). Minor variants were distributed across the entire genome (Figure 1C). We observed that some positions in the M, G, F, M2-1 and L genes showed higher frequencies relative to the rest of the genes. The NS1 and NS2 did not show minor variants occurring at high frequency. The NS1 region was difficult to sequence fully in majority of the samples (Figure1 B and C). The number of minor variants occurring at high frequencies were notable in G, M2-1 and M2-2 gene segments in addition to the L-gene (Figure 1B). The overall distribution of minority variants across the genome and from all the analysed samples is illustrated in Figure 1C. The uneven coverage was largely due to the amplicon-based amplification method of the genome.

**Figure 1:**
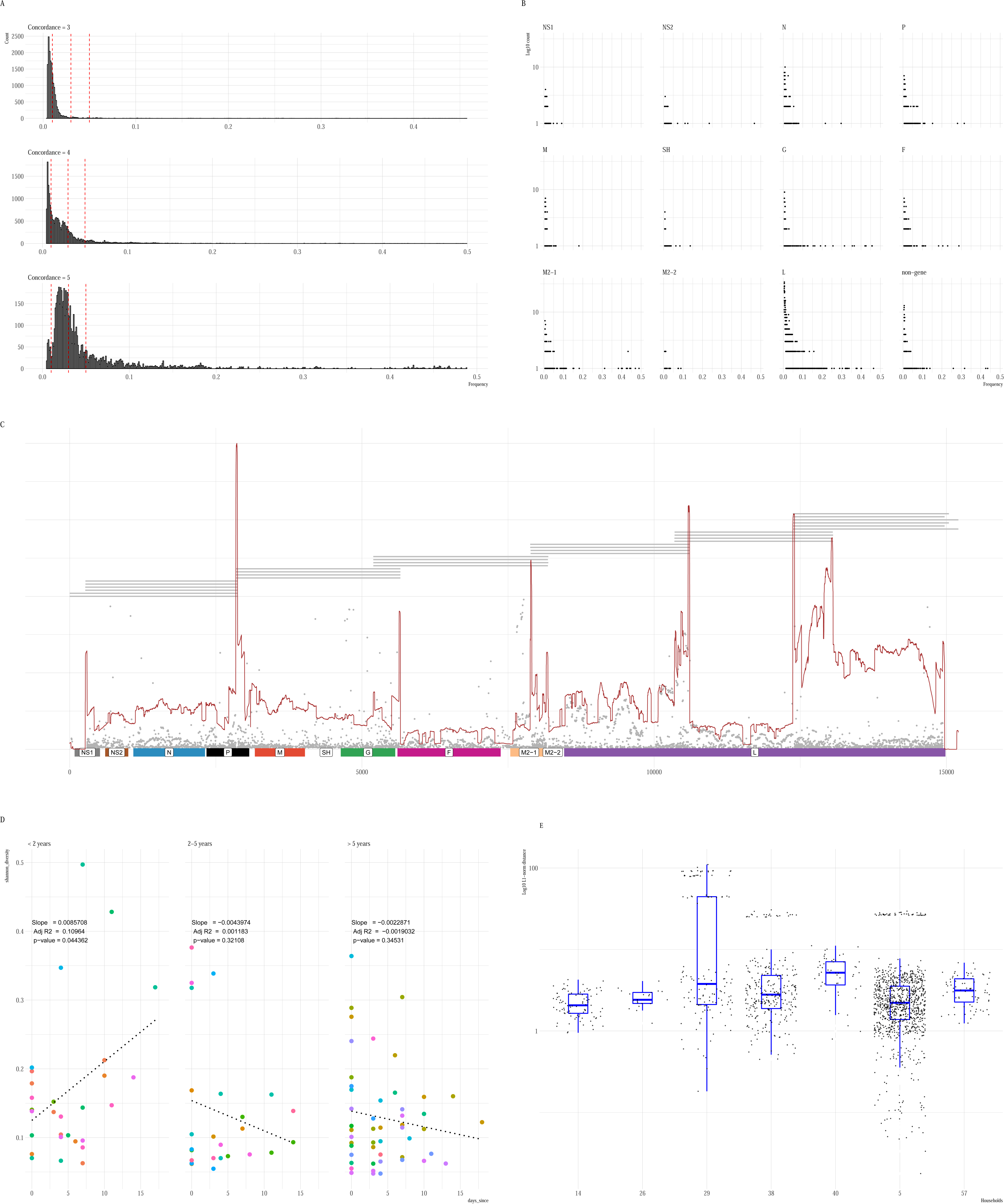
Summary of minority variants diversity of RSV-A from samples collected from a community epidemiological study. **A)** The frequency of minority variants stratified by concordance (3, 4, and 5). Concordance was defined as the agreement by different variant callers in identifying a variant position. The dashed vertical lines indicate the 5%, 3% and 1% error thresholds that are often used as arbitrary conservative cut-offs to distinguish between actual variants and errors. **B)** Plots showing the distribution and frequency of minority variants in both gene-coding and non-gene coding RSV regions. **(C)** The genomic location (x-axis) and frequency distribution (y-axis) of identified minority variants in the RSV A genome for xx samples. Each dot represents a minor variant at frequency below 0.5. The red line represents the read coverage shown using a normalized median value. The layout of the multiple amplicons used to amplify each genome are shown by the grey segments each of which corresponds to a primer-pair and the respective amplification target region. The 11 RSV gene coding regions are shown along the bottom panel. **D)** Scatter plots showing the relationship between within host diversity (Shannon’s diversity index) and the number of days since first positive sequenced sample shown for individuals of different age groups. The dotted line shows a linear fit of the observations for individuals in each age group. **E**) Boxplots showing the pairwise L1-norm genetic distances between each pair of individuals in each household.

We were interested in understanding the pattern of within host diversity among individuals in different age categories (less than 2 years, 2 to 5 years and above 5 years). We explored correlation between mean diversity (Shannon entropy) per sample against time since first positive sample by fitting a linear model between observed mean diversity and days since the correction of the sample. Overall there was no relationship between the diversity and days since first sample. We fitted a second model to account for age structure and we observed a positive linear relationship between mean Shannon’s entropy and time in samples collected from younger individuals (< 2 years) (Figure 1D).

Results from a recent study (Agoti et al. 2017) and that utilised the same dataset, showed household level clustering of consensus sequences. To test whether within host minority variant diversity was useful for describing within and between house diversity, we carried out a pairwise diversity assessment using the L1-norm distance between sample pairs (Figure 1E). We estimated the difference in within-household and between-household mean using a linear model to estimate the difference in mean denoted as ß1 between the within-household (µ1) and between-household (µ2) groups (Table 1).

**Table 1:** A table showing the estimated difference in mean genetic differences in samples from individuals within the same household vs individuals from different households. ß1 denotes the difference in mean between the two groups (within-household (µ1) and between-household (µ2).

| Term | Estimate | HDI (lower) | HDI (upper) |
| --- | --- | --- | --- |
| Within household mean ($\mu_1$) | 2.46 | 2.39 | 2.53 |
| Between household mean($\mu_2$) | 2.92 | 2.76 | 3.6 |
| Difference in mean($\beta_1$) | 0.46 | 0.37 | 0.54 |
| Within household variation ($\sigma_1$) | 0.08 | 0.02 | 0.15 |
| Normality parameter ($v$) | 1.04 | 1.01 | 1.08 |

Figure 2 shows the patterns of diversity among minority variant positions that were shared across samples collected from the same individuals. The minority variant frequencies from a few of the positions increased over time (Figure 2) while others decreased. Overall, they present a stochastic picture of diversity that is difficult to tease out given the number of samples versus the number of potential confounders.

**Figure 2:**
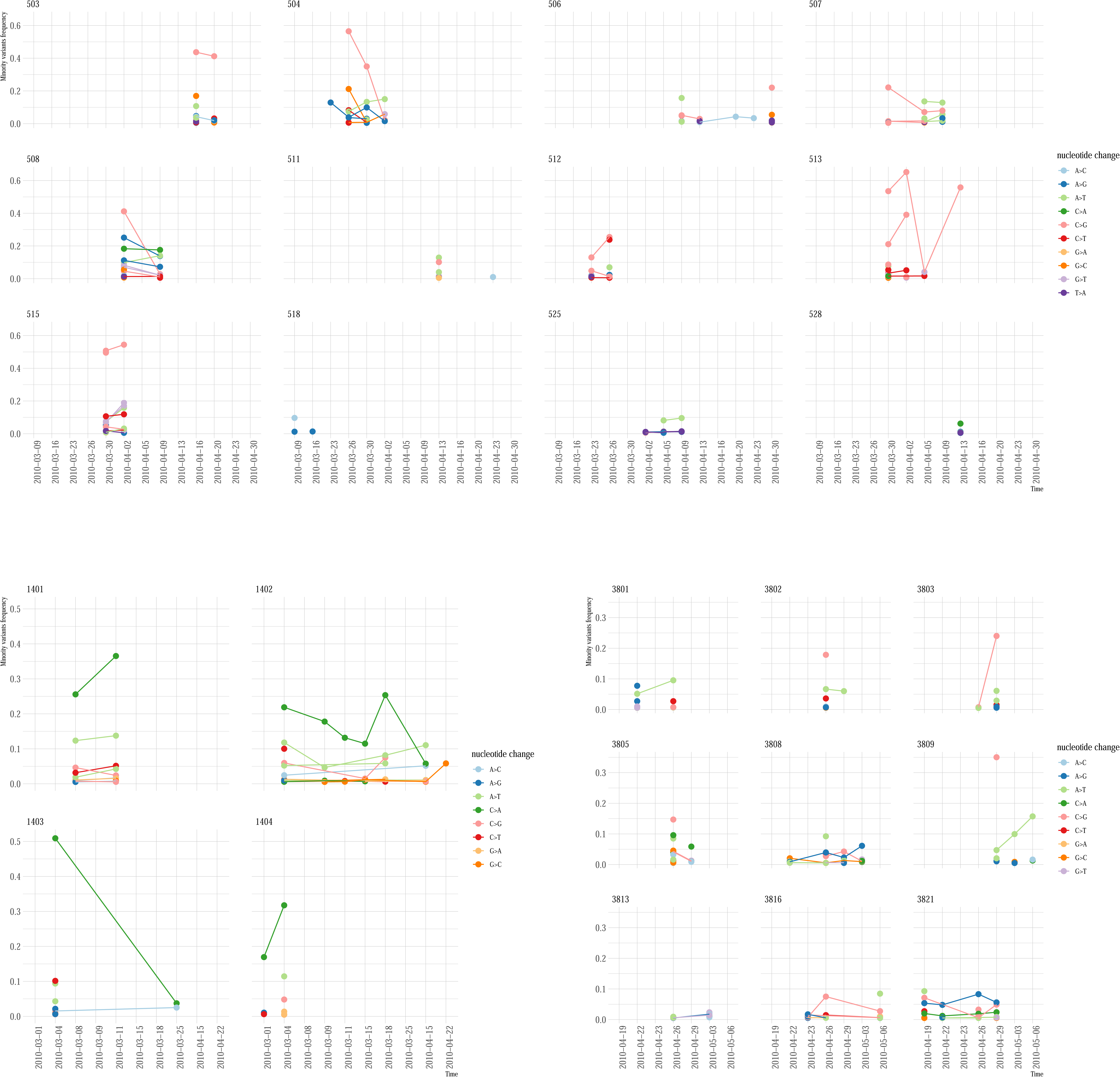
Summary of within host temporal fluctuations in the minority variant frequencies. Each panel shows variants that were detected across samples collected from the same individuals over time (x-axis). The y-axis shows the fluctuation in frequencies amongst these variants and colour denotes the type of nucleotide substitution.

### Application of shared minority variants to infer transmission patterns of RSV A

To evaluate the potential application of shared RSV-A diversity in providing additional resolution on who infects whom, we utilised shared variants from samples collected from individuals from three households. We restricted our analysis to the set of samples that comprised observable epidemiological clusters (Figures 3, 4 and supplementary figure 4). These cut-offs excluded apparent outliers according to an empirical distribution (Worby, Lipsitch, and Hanage 2017). Overall the analysis supports inference drawn from the consensus whole genome sequence phylogenetic tree, adds new information and at times provides contradictory insights from the consensus phylogenetic tree. Connected components from the minor variant networks were generally consistent with information from the phylogenetic tree (Figure 4C, 5C and 6C). To disentangle these data and information, we explored our observations on a case by case basis across three households (household 38 (HH38), household 14 (HH14) and household 5 (HH05)) that had variable membership and epidemiological dynamics.

**Figure 3:**
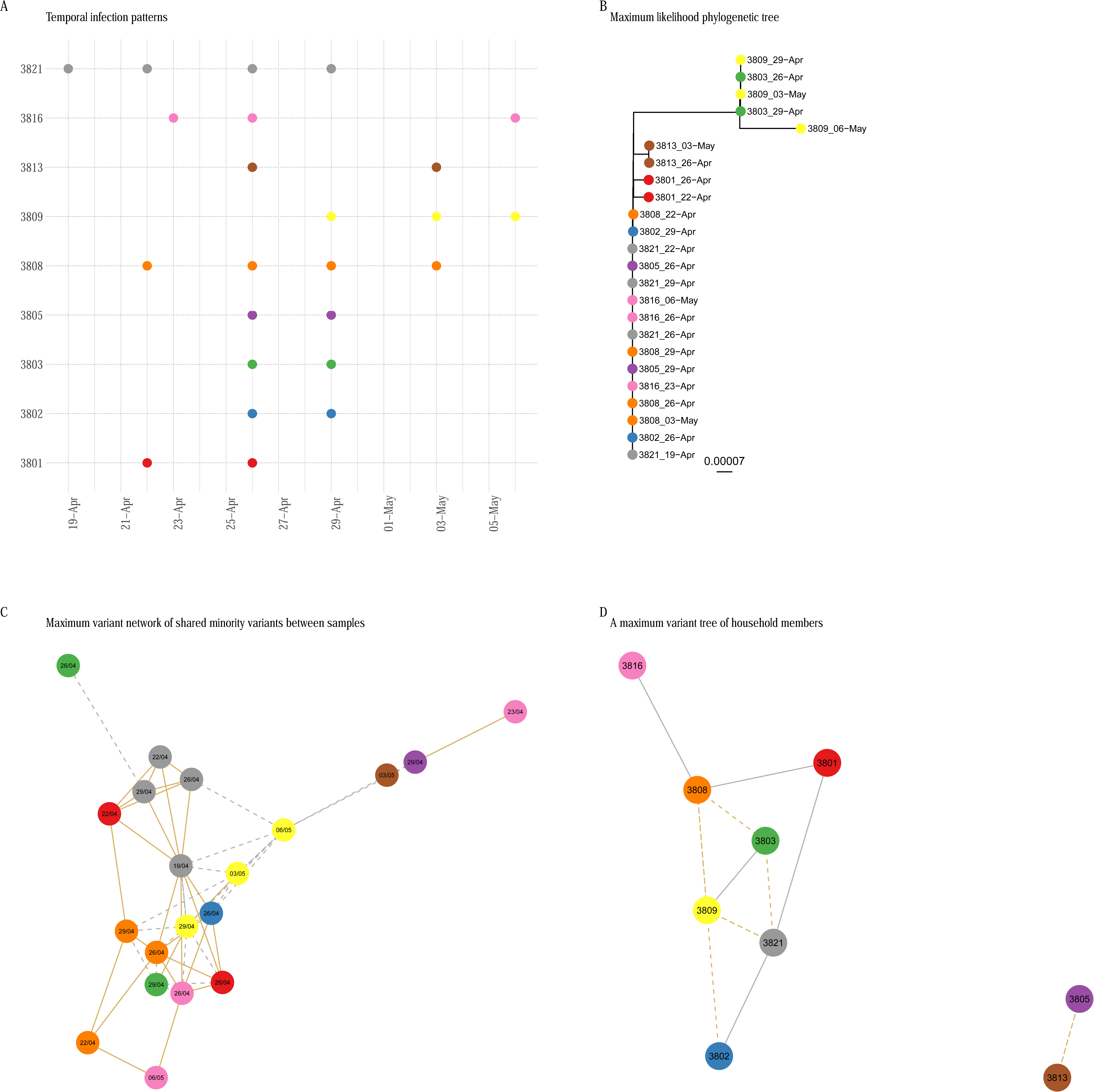
Putative transmission patterns in household 38 based on shared minority variants. **(A)** The RSV-A temporal infection pattern in household 38 (9 members) from the household study. Each filled circle represents a sequenced RSV-A positive sample (n=24) and colour denotes samples collected from the same individual. **(B)** A maximum likelihood phylogenetic tree created from an alignment of whole genome consensus sequences from each successfully sequenced sample. **(C)** A non-directed maximum variant transmission network drawn using shared variants from whole genome sequenced samples. Samples (vertices) are connected if they shared minority variants. Samples are connected by a dashed edge if their pairwise consensus sequences are separated by more than two SNPs **(D)** An individual based minority variant maximum weighted transmission tree. Here each node represents an individual from the household. Solid lines connect individuals who shared one or more samples with identical consensus sequences.

**Figure 4:**
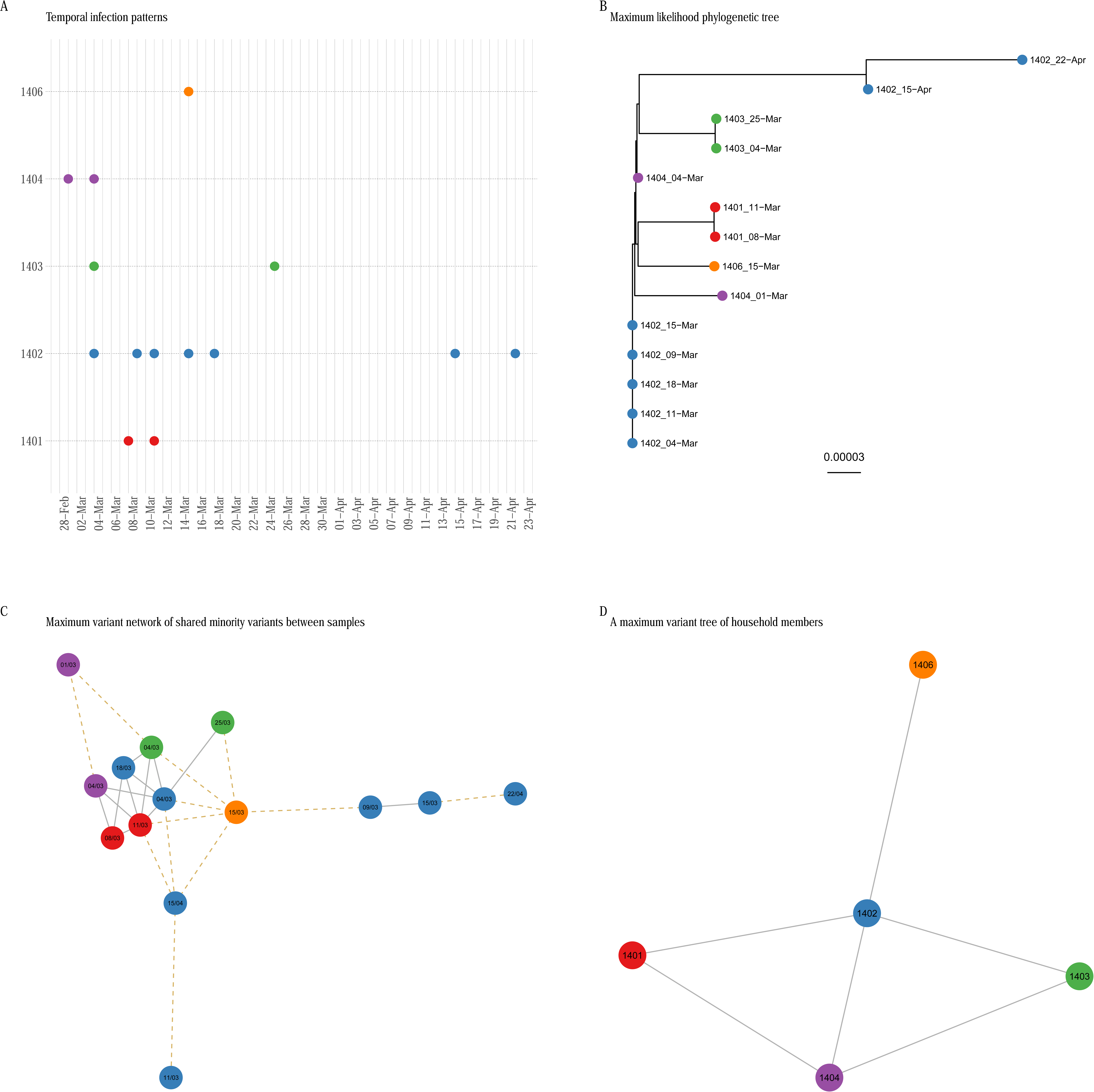
Putative transmission patterns in household 14 based on shared minority variants. **(A)** Epidemiological infection patterns in 5 members of household 14. The y-axis represents individuals and the x-axis represent time. Each filled circle represents a sequenced RSV-A positive sample (n=14). Samples collected from the same individual are denoted by the same colour. **(B)** A maximum likelihood phylogenetic tree created from an alignment of whole genome consensus sequences from each RSV-A sample collected from household 14. **(C)** A non-directed maximum variant transmission network drawn using shared variants from whole genome sequenced samples in A that met a predefined criterion. **(D)** An individual based minority variant maximum weighted transmission tree. Each node represents an individual from the household. An edge represents shared minority variants across all samples from the individual. The dashed lines indicate samples whose whole genome consensus sequences were separated by more than two SNPs.

Data from household 38 (HH38) suggested that individual 3821 was the first to be infected and presumably may have been the source of infection in this household. From the whole genome consensus data alone, (Figure 3B) samples collected from individual 3821 were identical to eleven samples collected from four individuals in the same household. Additional nine samples had more than one nucleotide difference collected from individuals 3801, 3803, 3809 and 3813. Sequences from samples collected from individual 3803 and 3809 formed a separate phylogenetic cluster. The minor variants diversity among members of this household was shared more frequently among samples with the same consensus sequence and presumably the same genetic background compared to samples that differed with one or more nucleotide changes at the consensus level. The earliest sample (19/4) collected from individual 3821 shared minor variants with samples collected from different individuals of related genetic background (same consensus sequence) in addition to samples collected later from the same individual (Figure 3C). The pattern of shared variants during the course of this outbreak, suggests that individual 3821 may have been a source of infection for individual 3801, 3802, 3803 and 3809. For example, we can deduce that the infant (3801) could have been infected by individual 3821 or 3808, but individual 3808’s first positive sample was collected on the same date as 3801, and therefore unlikely to have infected 3801. This assertion is supported by shared minor variants between the samples collected from these individuals. Contrary to phylogenetic inference, individual 3805 and 3813 appear to separate from the main minor variant sharing network component (Figure 3D) and they do not share minor variants with samples collected from other individuals. samples from individuals 3805 and 3813 clustered distinctly based on the pairwise genetic distance (L1-norm), which suggests that these samples were a separate introduction.

We explored if the minority variant information provided additional resolution on who infects whom amongst members of household 14 (HH14). HH14 was made up of five members (figure 4). The RSV-A positive samples could be considered as an epidemiological cluster (figure 4A). Individual 1406 was the oldest person in the household and individual 1401 was the infant. The phylogenetic information from whole genome consensus sequences suggested little genetic differences among these samples. Similar to previous observation from HH38, within host differences and their contribution to transmission is difficult to tease out based on the phylogenetic tree alone. While samples collected over consecutive days from individual 1402 were similar at the consensus level other than those collected on 15^th^ and 22^nd^ of April, the pattern of within host variation suggests that these samples shared substantial minor variants with each other (Figure 5). From figure 5A it appears that individual 1404 was the index case and presumably brought the infection to this household. This can be confirmed from the phylogenetic consensus tree.

Household 5 (HH5) contained a large number of infected individuals (n=19) and not all of all the collected samples (supplementary figure 3A) yielded sufficient sequence data to assemble whole genome consensus sequences (supplementary figure 3B). In addition, it was difficult to describe what comprises an epidemiological cluster from the temporal infection patterns alone. Whole genome phylogenetic tree showed considerable diversity between samples. However, there was a considerable number of shared variants between samples (supplementary figure 3C and 3D). In this case interpreting the patterns transmission from minority variant data is a lot more challenging owing to insufficient genomic data for example only 3 of 5 samples from individual 518 and 506 respectively were available. Only a single positive sample was collected from individual 514 and which appears to present a potential separate introduction. Furthermore, only a single sample was sequenced from the infant (individual 502). Despite these challenges, the sample from individual 502 comprised the same consensus sequence with those collected from individuals 504,506,507,512,515, 518,528,529 and showed a number of shared minority variants. From the pattern of shared minority variants, the data suggests that the infant 502 was likely infected by the eleven-year-old individual 518 or by the two-year-old individual 508.

## Discussion

Developing an effective control program against RSV transmission depends on ability to focus the control measure to a precise segment of the community or individuals who are at risk or those that are key links in the transmission path. Such an intervention requires obtaining information on host contact structure and mobility, in addition to the genetic diversity of the rapidly spreading virus. We are interested in uncovering the factors or parameters that are associated with successful spread and transmission of respiratory illness and consequently it is imperative to know who infects whom within the household. Presumably we can then go back and ask questions on the covariates that are associated with individuals who are connected by a direct transmission. This kind of information is difficult to obtain from epidemiological case detection alone, and hence utilization of virus genomic data has become routine. Several studies have used shared minority variants to infer host to host transmission patterns (Stack et al. 2013; Morelli et al. 2012; Poon et al. 2016; Worby, Lipsitch, and Hanage 2017; Paterson et al. 2015).

In this analysis we set out to assess the utility of shared low frequency diversity in understanding RSV transmission pathways to a unique epidemiological dataset (Munywoki et al. 2014) collected during an RSV community outbreak and which we have described previously (Agoti et al. 2017). Application of minority variants in epidemiological studies has become possible partly because deep sequencing provides an opportunity to obtain and examine virus diversity and evolutionary dynamics during an outbreak at a far greater resolution than temporal epidemiological case detection. Recent methods have applied loss and fixation of minor variant types within host to inform on pathogen population dynamics during viral shedding, and shared variants between different hosts to infer transmission chains (Poon et al. 2016; Worby, Lipsitch, and Hanage 2017; Stack et al. 2013; Paterson et al. 2015) and to estimate the size of the transmission inoculum (Sobel Leonard et al. 2017; McCrone et al. 2017). In these studies, it is assumed that the presence of the same nucleotide variant in two different hosts indicates that either both hosts are linked by a transmission event and that two or more variants survived the transmission bottleneck. It could also be interpreted that the shared mutation arose independently within each host or that contamination or sequencing error resulted in the apparent shared variation. By considering only variants on the same genetic background and applying a conservative variant frequency threshold, the remaining shared variants could be indicative of direct transmission (Worby, Lipsitch, and Hanage 2017). Simulation testing showed that even with unsampled cases and homoplasy this method was robust at identifying direct transmission where it exists.

In a previous study and using the same set of samples, consensus RSV virus sequences and a phylogenetic method were used to describe transmission patterns across household 14 and household 38. In general the study provided a description of virus movement that was consistent with the household membership (Agoti et al. 2017). However, it has been shown that inference of transmission chains from phylogenetic tree is not always direct or straightforward (Pybus and Rambaut 2009; Romero-Severson et al. 2014) and that the transmission tree topology can differ remarkably from the phylogeny (Kenah et al. 2016). Here we wonder if including minority variant data increases the resolution provided by the consensus sequence data. We did this by quantifying the minority variant population of each RSV-A positive sample and explored the application of shared within-host diversity in providing additional resolution to understand who infects whom. We profiled the distribution of minority variants across the RSV genome and showed that intra-host diversity is non-uniform. The data suggests that minority variants in RSV genomes occurs at low frequency (Figure 1B and C). The G-gene, the M1-2 and the L gene including the non-coding regions of the genome appear to be more amiable to low frequency variation. Furthermore, pairwise genetic diversity from samples collected from the same household was lower compared to individuals from different households (Figure 1E).

To infer putative transmission pathways, we argue that that RSV diversity is transmissible and that in order for variants to be useful to explore transmission routes, the bottleneck cannot be strict. Evidence that it may not be strict is indirectly supported by studies that have showed that RSV is very contagious and spread via contact and large droplets over short distance (Hall 1983) and that the virus can persist on surfaces for up to 1 hour (Hall, Douglas, and Geiman 1980). Our data provides evidence for this postulate by the observation that we detected shared variants between samples (Figure 3 & 4, supplementary figures 3) even after applying a stringent and conservative variant filtering criterion. The analysis was largely focused on samples that constituted what appeared as epidemiological clusters. This was important to avoid shared variants arising from homoplasy given that mutations can arise randomly in sequences from different genetic backgrounds.

Using minority variant data provided an additional layer of data and insight. For example, we observed that samples with identical consensus sequences provide a dynamic picture of within host diversity. In terms of transmission, some samples shared more variants (Figure 3C, supplementary figures 4C, 5C) comprising of cliques of shared variants. This implies that intensive sampling provides more information relative to one off sampling and that within host variation is important in understanding transmission for rapidly evolving pathogens. We observed both agreement and disagreement between the consensus phylogenetic transmission tree and the minor variant derived transmission trees. These patterns varied from household to household. Large households (supplementary figure 5) provide a particular challenge because there was a lot of heterogeneity and possible multiple introductions which are difficult to tease out.

Our approach is not without limitations, first, quantification of minority variant population is dependent on the quality of the clinical sample, the errors introduced by sample handling, the sequencing process and the minority variant detection algorithm. We utilised a concordance based approach for detecting and quantifying minor variants (Said Mohammed et al. 2018) to mitigated against these sources of error. Secondly, defining an appropriate minority variant frequency threshold and establishing informative threshold for shared minority is a challenge because this does not fully address the fundamental question in that, we do not know the sources for each infection and the genetic patterns associated with each transmission when presented with data from a natural epidemic. For that, we require deliberate transmissions, where the source is known, so that we can see what the patterns look like at the genetic level. Therefore, it is possible that infections within the households are from outside the household so that the observed patterns are due to having shared sources (which are outside the households) rather than direct transmission. Finally, we assume that diversity is transmissible and thereby we apply genetic information to inform transmission trees. The essence is to have a probability that two infections are linked using genomic data in addition to the probability provided by the epidemiological information. However, these two probabilities do not correlate perfectly, and further work is required to test this assumption.

## Conclusion

In conclusion, we have described and made use of shared minority variants to infer within household transmission patterns of RSV-A. Minority variants in RSV occur at very low frequencies. We assessed the utility of minority variants in providing additional resolution in addition to phylogenetic trees, derived from RSV-A whole genome consensus sequences. We showed that RSV-A infections are characterised by low frequency variants and although they provide additional information where consensus sequences are identical, and as a consequence make the phylogenetic signal unhelpful in identifying transmission, relying on minor variants alone is insufficient for inferring transmission pathways for RSV household infections. Not all low frequency nucleotide changes over time are useful because they may not have arisen in the course of a transmission event. Single sample sequencing is inadequate to capture the proportion of minority variants that are associated with transmission. Additional research is required to develop a detailed probabilistic framework using minor variants data and describe not only who acquires infection from whom but also the broader questions such as the effect of generation time, mutation rates and selection on transmission and that may provide us with quantifiable parameters that characterize transmission of respiratory infections.

## Acknowledgments

We are thankful to Virus Epidemiology and Control Group in Kilifi, the field and clinical staff who collected, processed the samples in addition to the guardians and parents of the children who participated in the study. This study was published with the permission of the director of KEMRI.

## Grant information

The work was funded by the Wellcome Trust (Wellcome Trust Grant references: 077092, 090853, 100542 and 102975) in addition to the DELTAS Africa Initiative [DEL-15-003]. The DELTAS Africa Initiative is an independent funding scheme of the African Academy of Sciences (AAS)’s Alliance for Accelerating Excellence in Science in Africa (AESA) and supported by the New Partnership for Africa’s Development Planning and Coordinating Agency (NEPAD Agency) with funding from the Wellcome Trust [107769/Z/10/Z] and the UK government. The views expressed in this publication are those of the author(s) and not necessarily those of AAS, NEPAD Agency, Wellcome Trust or the UK government.

**Supplementary Figure 1:**
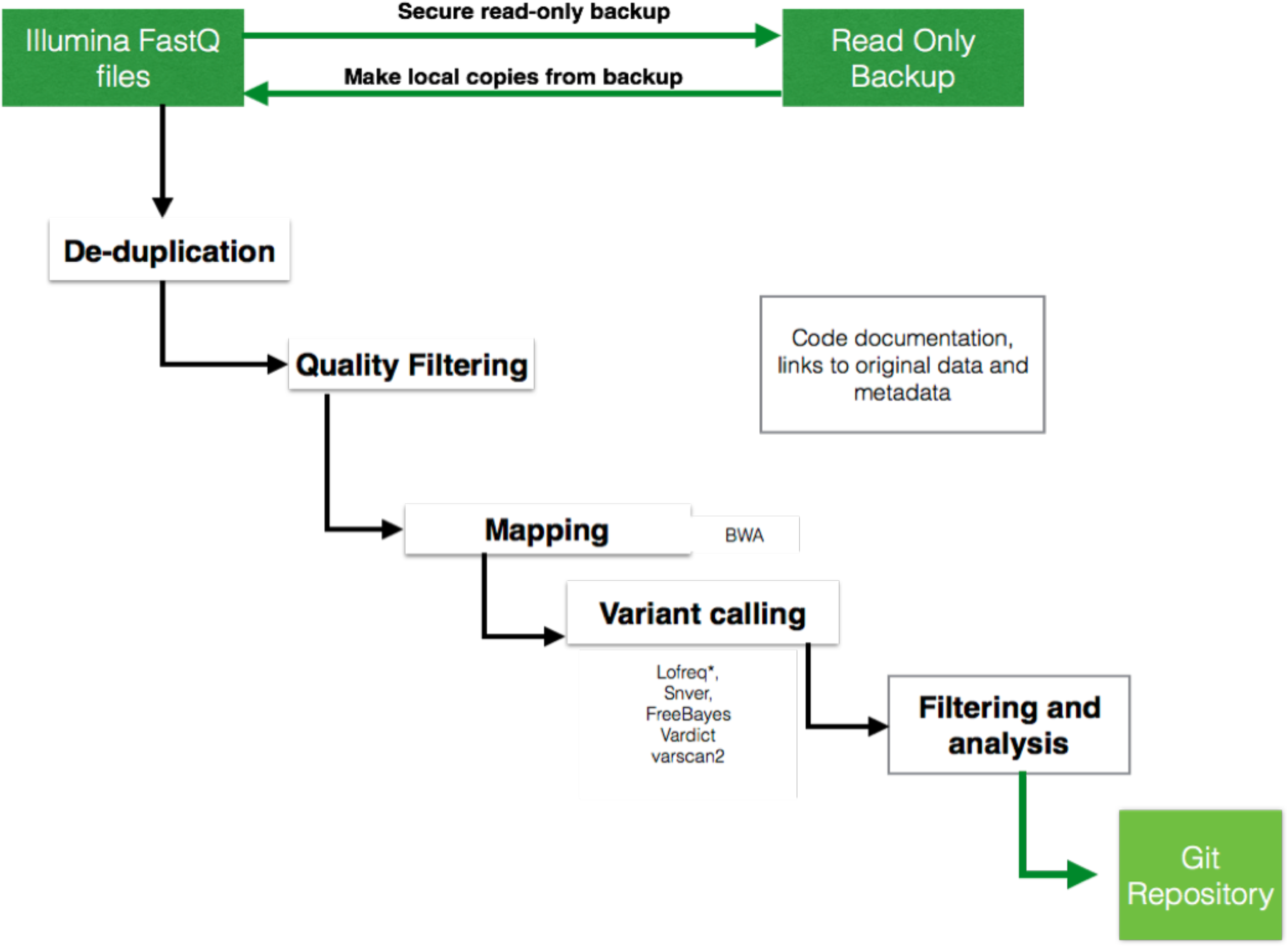
A summary of the bioinformatics workflow. A schematic diagram showing an overview of the bioinformatics workflow used to process the sequence data and call the minority variants from short-read paired end Fastq files. Low quality and duplicated reads were discarded, and the remaining reads were trimming at the 3’ and 5’ ends such that only positions with base quality scores (PHRED) >30 were retained. Minority variants were called using five published minority variant callers and each variant position was assigned a concordance value that corresponds to the number of minority variants callers that identified the variant position.

**Supplementary Figure 2:**
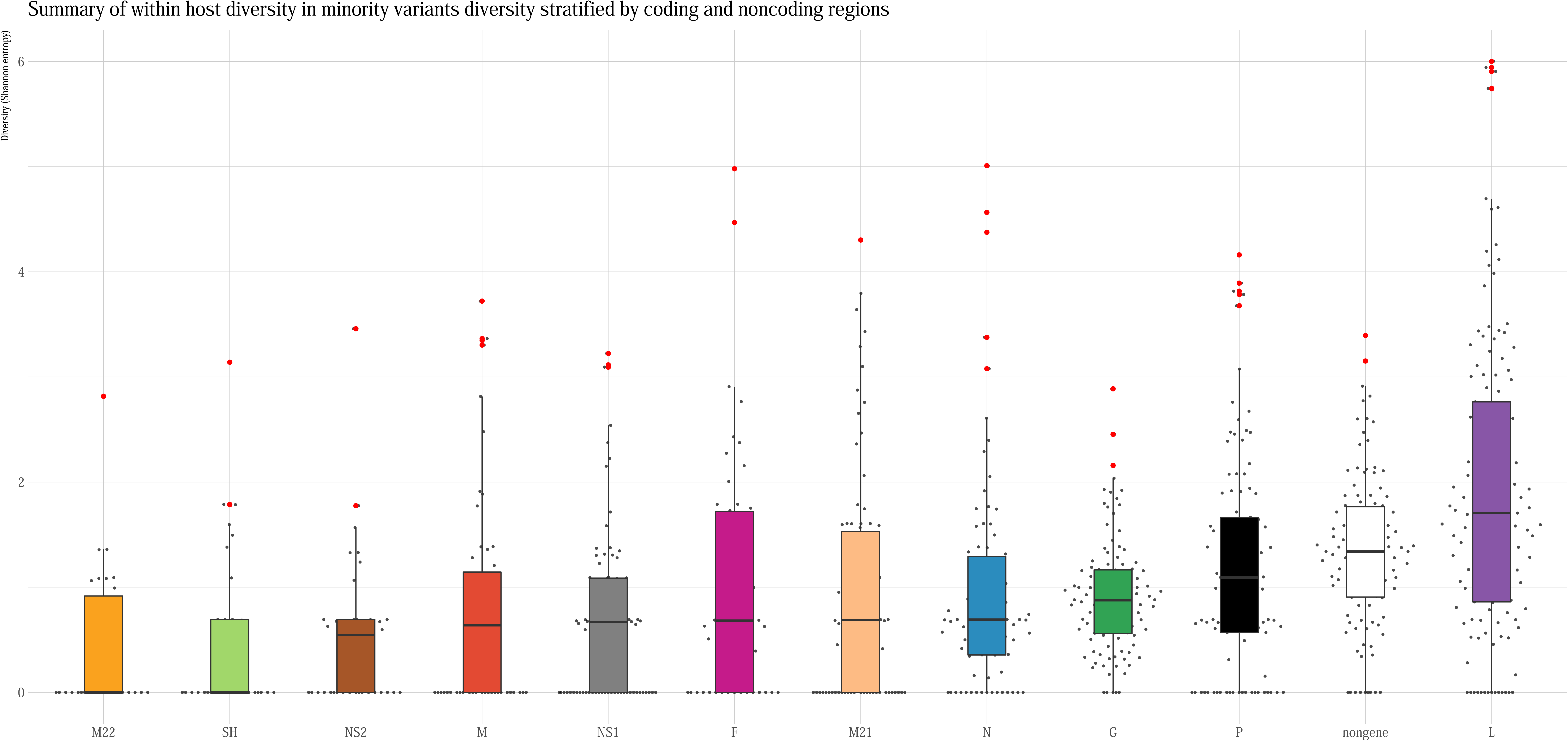
A summary of per gene within-host minority variants diversity. A plot of the median RSV-A Shannon’s entropy sorted from left to right.

**Supplementary Figure 3:**
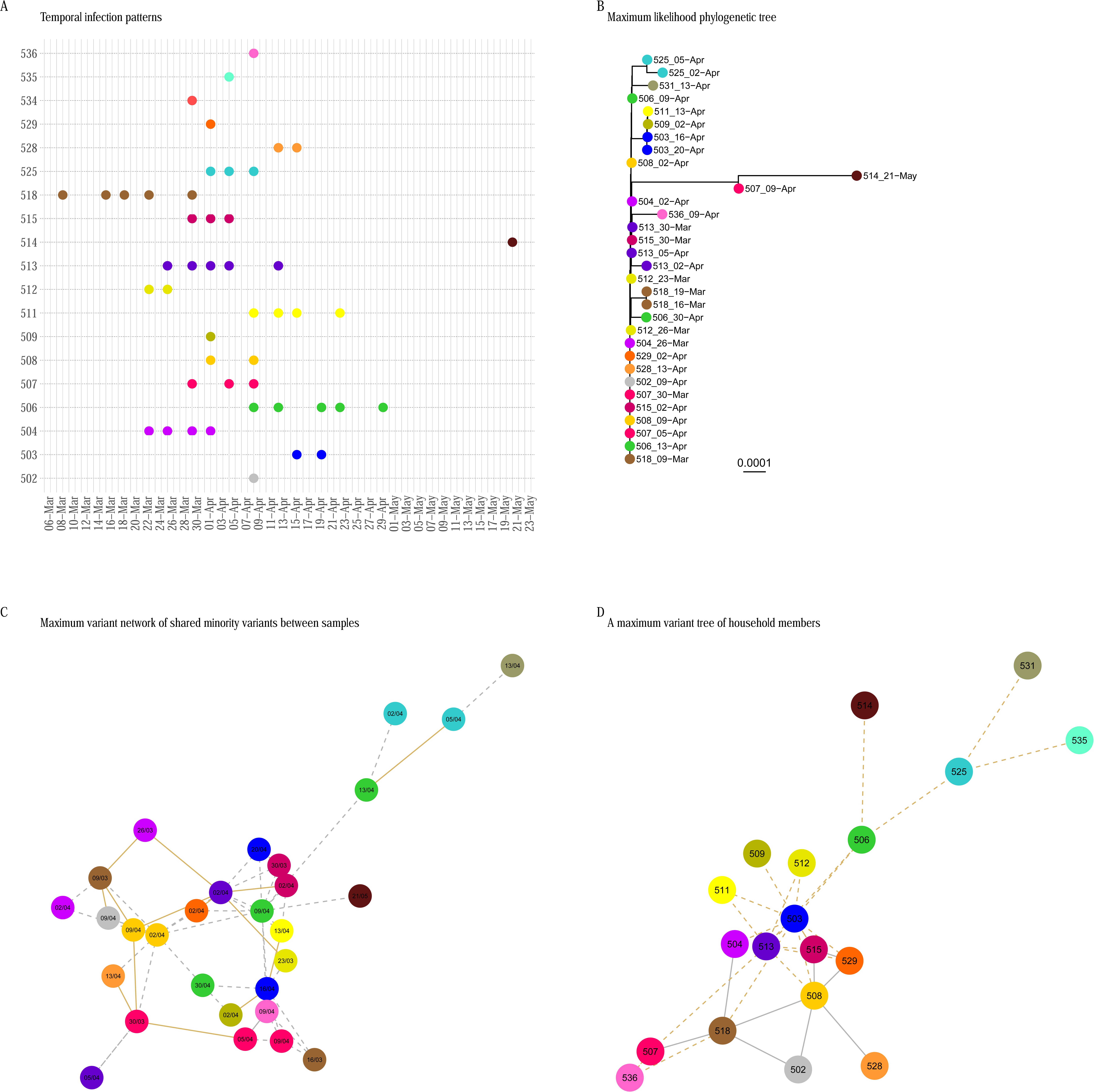
Putative transmission patterns in household 5 based on shared minority variants. **(A)** The epidemiological infection profiles from 19 members of household 5. Each filled circle represents a sequenced RSV-A positive sample for a total of 45 samples. Colour represents samples collected from the same individual. (B) A maximum likelihood phylogenetic tree from consensus whole genome sequence. The colours denote samples from the same individual. **(C)** A non-directed maximum variant transmission network drawn using shared variants from whole genome sequenced samples in A that met a predefined criterion. **(D)** An individual based minority variant maximum weighted transmission tree. Individuals (vertices) are connected by a solid line (edge) if one or more of their samples had an identical consensus sequence in common. In both **C** and **D** panels an edge represents shared minority variants across all samples from the individual. The dashed lines indicate samples whose consensus sequences were separated by more than two SNPs.

